# Frequency of disturbance alters diversity, function, and underlying assembly mechanisms of complex bacterial communities

**DOI:** 10.1101/313585

**Authors:** Ezequiel Santillan, Hari Seshan, Florentin Constancias, Daniela I. Drautz-Moses, Stefan Wuertz

**Author notes:** Correspondence to: Stefan Wuertz. Brown and Caldwell, 9665 Chesapeake Drive, Suite 201, San Diego CA 92123, U.S.A.

## Abstract

Disturbance is known to affect ecosystem structure, but predicting its outcomes remains elusive. Similarly, community diversity is believed to relate to ecosystem functions, yet the underlying mechanisms are poorly understood. Here, we tested the effect of disturbance on the structure, diversity, and ecosystem function of complex microbial communities within an engineered system. We carried out a microcosm experiment where activated sludge bioreactors were subjected to a range of disturbances in the form of a toxic pollutant, tracking changes in ecosystem function. Microbial communities were assessed by combining distance-based methods, general linear multivariate models, *α*-diversity indices, and null model analyses on metagenomics and 16S rRNA gene amplicon data. A stronger temporal decrease in *α*-diversity at the extreme, undisturbed and press-disturbed, ends of the disturbance range led to a hump-backed pattern, with the highest diversity found at intermediate levels of disturbance. Undisturbed and press-disturbed levels displayed the highest community and functional similarity across replicates, suggesting deterministic processes were dominating. The opposite was observed amongst intermediately disturbed levels, indicating stronger stochastic assembly mechanisms. Tradeoffs were observed in community function between organic carbon removal and both nitrification and biomass productivity, as well as between diversity and these functions. Hence, not every ecosystem function was favoured by higher community diversity. Our results show that the assessment of changes in diversity, along with the underlying stochastic-niche assembly processes, is essential to understanding the impact of disturbance in complex microbial communities.

**Importance:** Microbes drive the Earth’s biogeochemical cycles, yet how they respond to perturbations like anthropogenic pollutants is poorly understood. As human impact continues to increase worldwide, foreseeing how disturbances will affect microbial communities and the ecosystem services they provide is key for ecosystem management and conservation efforts. Employing laboratory-scale wastewater treatment bioreactors, this study shows that changes in community diversity accompany variations in the underlying deterministic-stochastic assembly mechanisms. Disturbances could promote stochastic community structuring, which despite harboring higher diversity could lead to variable overall function, possibly explaining why after similar perturbations the process outcome differs. A conceptual framework, termed the ‘intermediate stochasticity hypothesis’ is proposed to theoretically predict bacterial community shifts in diversity and ecosystem function, given a range of possible disturbance types, in a well-replicated time-series experiment. Our findings are relevant for managing complex microbial systems, which could display similar responses to disturbance, like oceans, soils or the human gut.

## Introduction

Understanding what drives patterns of community succession and structure remains a central goal in ecology (1, 2) and microbial ecology (3), especially since community diversity and assembly are thought to regulate ecosystem function (4, 5). The factors influencing the balance between mechanisms of community assembly are under debate and require studies across a range of ecosystems (6, 7). Assembly processes can be either stochastic, assuming that all species have equal fitness and that changes in structure arise from random events of ecological drift (8), or deterministic, when communities form as a result of niche diversity shaped by abiotic and biotic factors (9). Deterministic and stochastic assembly dynamics have been proposed to simultaneously act in driving assembly patterns observed in nature (10-14). This has stimulated scientific discourse including modelling of experimental data (15-18) and both observational and manipulative experimentation in a variety of ecosystems, like deserts on a global scale (19), groundwater (6), subsurface environments (2, 20, 21), soil plant-fungi associations (22), rock pools (23), water ponds (24), and sludge bioreactors (7, 17, 25). These prior studies emphasized the need to understand what governs the relative balance between stochastic and deterministic processes and what conditions would lead to stochastic processes overwhelming deterministic processes, particularly under disturbance (21). To investigate their roles well-replicated time series experiments are needed (6, 25).

Disturbance is defined in ecology as an event that physically inhibits, injures or kills some individuals in a community, creating opportunities for other individuals to grow or reproduce (26). Under disturbance, organisms may benefit from being able to grow, reproduce, and interact with other members of the community in the absence of specific familiarity with the environment. It is deemed a main factor influencing variations in species diversity (27) and structuring of ecosystems (28, 29), but a clear understanding of its outcomes is lacking (30). Particularly, the intermediate disturbance hypothesis (IDH) (31) predicts that diversity should peak at intermediate levels of disturbance due to trade-offs between species’ ability to compete, colonize ecological niches, and tolerate disturbance. The IDH has been influential in ecological theory, as well as in management and conservation (32), but its predictions do not always hold true (27, 33). For example, in soil and freshwater bacterial communities different patterns of diversity were observed with increasing disturbance frequency with biomass destruction (34) and removal (35) as disturbance type, respectively. Meanwhile, the effect of varying frequencies of non-destructive disturbances on bacterial diversity remains unknown. Furthermore, the IDH predicts a pattern but it is not a coexistence mechanism as it was originally purported to be (36). Hence, its relevance is being debated (37, 38) with multiple interpretations and simplicity as the main points of critique. To date, the mechanisms behind the observed patterns of diversity under disturbance remain to be elucidated (39, 40).

The objective of this work was to test the effect of disturbance on the bacterial community structure, diversity, and ecosystem function of a complex bacterial system, with emphasis on the underlying assembly mechanisms. We employed sequencing batch bioreactors inoculated with activated sludge from an urban wastewater treatment plant, in a laboratory microcosm setup with varying frequencies of augmentation with toxic 3- chloroaniline (3-CA) as disturbance. Chloroanilines are toxic and carcinogenic compounds and few bacteria encode the pathways to degrade 3-CA (41), which is also known to inhibit both organic carbon removal and nitrification in sludge reactors (42). Microcosm studies are useful models of natural systems (43), can be coupled with theory development to stimulate further research (44), and by permitting easier manipulation and replication can allow inference of causal relationships (45) and statistically significant results (46).

We analyzed changes in ecosystem function over time by measuring removal of organic carbon, ammonia, and 3-CA, as well as biomass. Changes in community structure were examined at different levels of resolution using a combination of metagenomics sequencing and 16S rRNA gene fingerprinting techniques. We also explored how diversity was related to function, focusing on tradeoffs. Furthermore, the role of stochasticity on community assembly was investigated by employing null model techniques from ecology.

## Results

### Overall community dynamics and differentiation of clusters

Bacterial community structure displayed temporal changes and varied between disturbance levels (Fig. 1). Constrained ordination showed a defined cluster separation with 0% misclassification error of the outermost levels L0 and L7 from the remaining intermediate levels L1-6 (Fig. 1A). Overall community structure differed over time with a dispersion effect after 14 days (Fig. 1B). Levels across disturbance and time factors showed significant differences (PERMANOVA P = 0.003, Table S1), with a non-significant interaction effect (P = 0.15). Disturbance was the factor responsible for the observed clustering (Fig. 1A), independent of dispersion heterogeneity (PERDIMSP P = 0.35).

**FIG 1.**
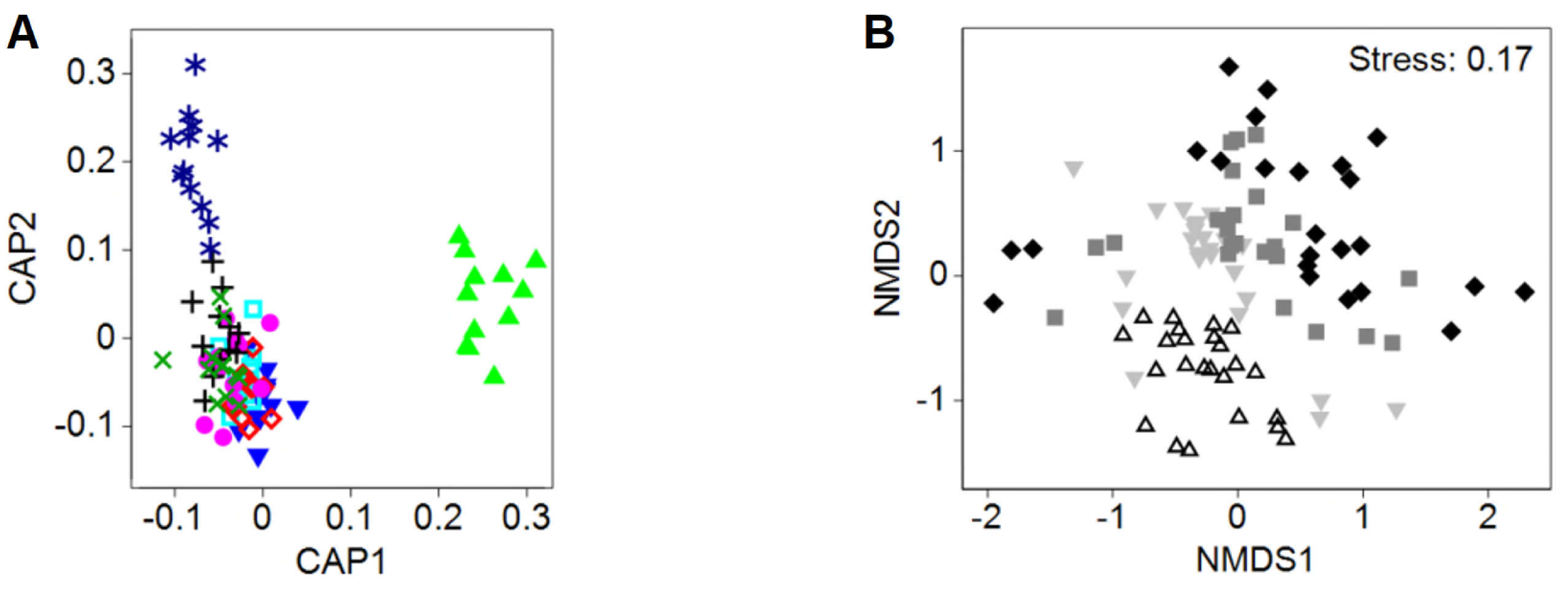
Ordination plots differentiating undisturbed (L0) and press-disturbed (L7) levels from intermediately disturbed levels (L1-6) and displaying temporal community dispersion effects for all time points. (A) Canonical Analysis of Principal coordinates (CAP, constrained ordination) plot, with disturbance levels as differentiation criteria, shows niche differentiation for L0 (CAP1 axis) and L7 (CAP2 axis). Disturbance levels: L0[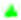], L1[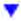], L2[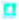], L3[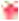], L4[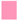], L5[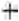], L6[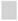], and L7[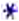]. (B) Non-metric Multidimensional Scaling (NMDS, unconstrained ordination) shows dispersion effect after a given number of days. Days: 14[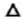], 21[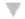], 28[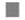], and 35[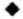].

### Ecosystem function dynamics and trade-offs

The undisturbed community (L0) was the only one with complete organic carbon removal and nitrification, while the press-disturbed community (L7) was the only one that could never nitrify and also had the lowest biomass (Fig. 3). Initially, reactors at the disturbed levels showed an inability to remove all of the 3-CA (with the exception of L1). Such lack of 3-CA degradation was accompanied by a reduction in organic carbon removal in the first three weeks (Fig. 2A, Fig. S3A-C), and a complete inhibition of nitrification with subsequent accumulation of ammonium (Fig. 2B, Fig. S3F-H). Removal of 3-CA recovered and was above 95% for all disturbed levels after 28 days (Fig. S3D), but COD removal was still not 100% despite complete 3-CA removal towards the end of the experiment (Figure 2C).

**FIG 2.**
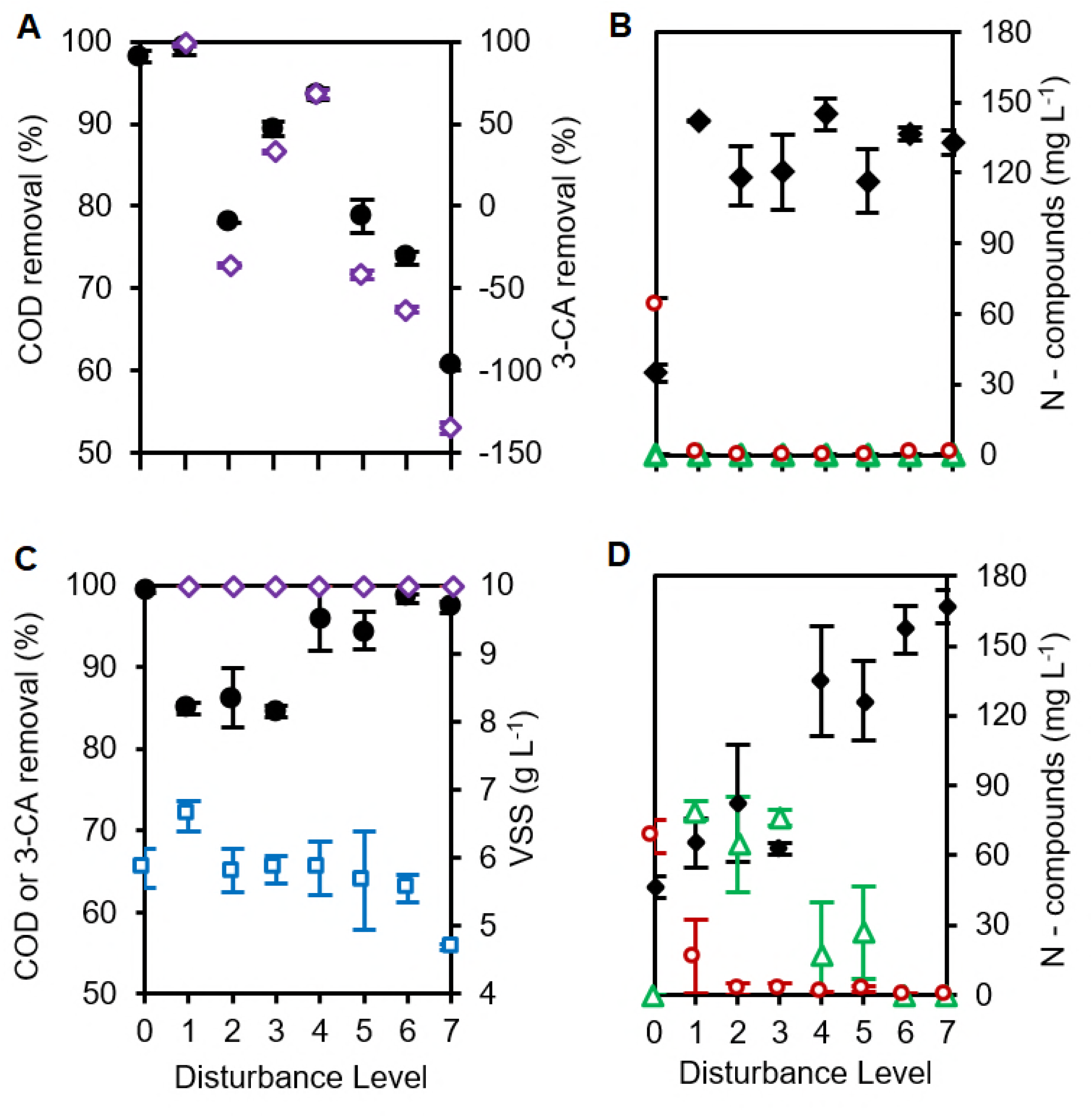
Process performance indicators across disturbance levels. Effects include temporal changes and tradeoffs in community function. (A,C) Percentage of organic carbon as chemical oxygen demand (COD, 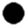) and 3-CA (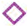) removal for all levels (negative values represent accumulation). (C) Biomass as volatile suspended solids. (VSS, 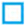). (B,D) Concentration of ammonium (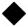), nitrite (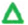), and nitrate (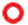) as nitrogen for all levels. Data are from days 7 (A-B) and 35 (C-D) of the study (for all time points sampled, see Fig. S2). Mean ± s.d. (n = 3) are shown. Undisturbed L0 replicates had consistent organic carbon removal and complete nitrification, whereas press-disturbed L7 never showed nitrification and had the lowest final biomass. Intermediate levels L1-6 displayed changing functionality with higher s.d. values that increased over time.

**FIG 3.**
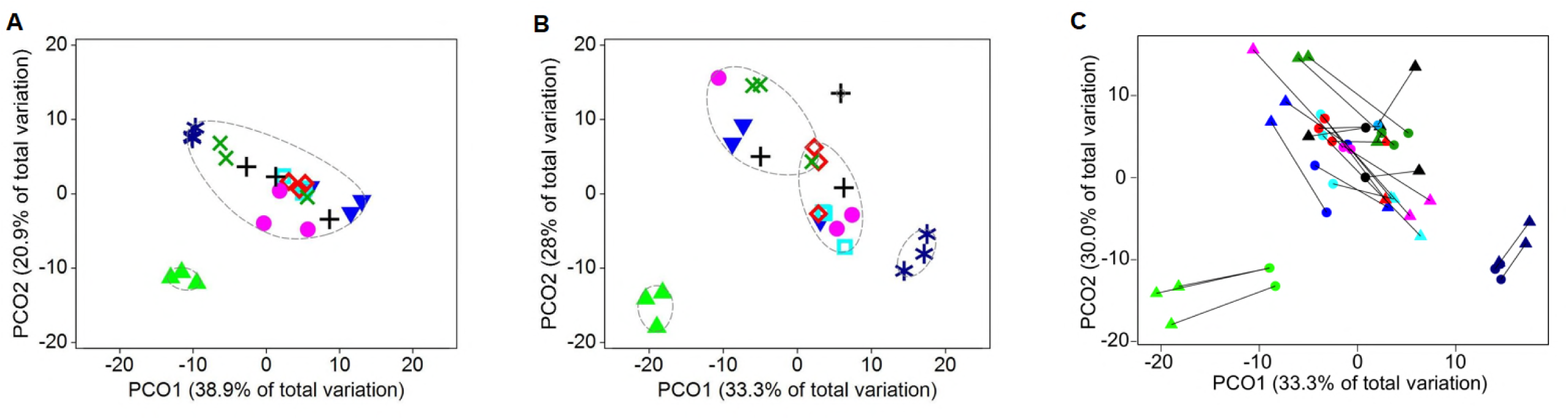
Community assembly as assessed by Principal Coordinates Analysis (PCO) plots for all disturbance levels on T-RFLP datasets on days (A) 14 and (B) 35 of the study. Ovals with dashed lines represent 80% similarity calculated by group average clustering. Disturbance levels: L0 [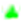], L1 [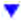], L2 [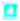], L3 [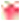], L4 [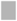], L5 [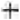], L6 [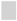], and L7 [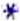]. (C) Procrustes analysis on PCO at day 35 comparing metagenomics (circles) and T-RFLP (triangles) datasets. Lines unite data points from the same reactor (n = 24). Same colour palette as for disturbance levels. Tests comparing both methods were statistically significant (Table S1). Intermediate treatments’ (L1-6) within-treatment dissimilarity increased with time. L0 and L7 clusters consistently displayed higher similarity after 14 days.

Nitrification was detected on day 21 for L1 and later for other disturbance levels, except for the press-disturbed L7. The dominant NO_X_ component was nitrite, but some nitrate was also produced (Fig. 2D, Fig. S2H-J). At the end of the study, there was a strong negative correlation between organic carbon removal and nitrification (Fig. S3A-B) in terms of nitrite production (r = −0.982, P = 0.003) and nitrate production (ρ = −0.697, P = 0.003). Biomass values on day 35 differed significantly among levels with the highest value at L1 and the lowest at L7 (Fig. 2C). There was a strong negative correlation between biomass and organic carbon removal (r = −0.418, P = 0.042), and positive between biomass and partial nitrification to nitrite (r = 0.488, P = 0.022) (Fig. S3C-E).

### Intermediate levels of disturbance displayed increased dissimilarity with time

To distinguish the effect of disturbance from temporal community dynamics (Fig. 1), community assembly was assessed at each time point by ordination analysis using PCO (Fig. 3A-B), NMDS, and CAP with cluster similarity analysis (Fig. S4). The combination of constrained and unconstrained ordination methods allowed differentiating location from dispersion effects in community assembly (47). L0 was consistently different in all ordination plots and L7 differed after 21 days, both with 0% misclassification error at all time points for CAP plots. Dispersion effects within intermediate levels were evident in the unconstrained ordination plots with higher differentiation of biological replicates after 35 days (Fig. 3B), coinciding with the production of nitrite and low levels of nitrate (Fig. 2D). Community differentiation was statistically significant from day 21 onwards as supported by multivariate tests (Table S1).

### Metagenomics community analysis validates observations from fingerprint dataset

**β**-diversity patterns observed from 16S rRNA gene amplicon T-RFLP data on day 35 were significantly similar to those from shotgun metagenomics data. A Mantel test on Bray-Curtis distance matrixes for both datasets (n = 24) yielded significant similarity (r = 0.73, P = 0.002). Procrustes tests of comparisons within ordination methods of PCO (Fig. 3C) and NMDS also yielded significant similarities for both datasets (P = 0.002, Table S1). Multivariate PERMANOVA tests on the metagenomics dataset produced statistically significant results, but with significant heteroscedasticity as shown by PERMDISP (Table S1). We resolved these mean-variance relationship concerns by running a General Linear Multivariate Models (GLMMs) test to fit the data to a negative binomial distribution. Both residuals vs fitted and mean-variance plots supported the choice of a negative binomial distribution for the regression model (Fig. S5). The analysis of deviance of the regression rejected the null hypothesis of no difference between communities at different disturbance levels, independent of heteroscedasticity (p = 0.0149).

### Higher α-diversity for intermediately disturbed treatments and its trade-offs with function

The observed patterns in *α*-diversity were time dependent, as diversity decreased over time with respect to the initial sludge inoculum (Fig. 4A). Such temporal decrease in diversity was higher at the extreme ends of the disturbance range, resulting in a parabolic pattern on day 35 (Fig. 4B-C). The final *α*-diversity pattern based on Hill number ^2^D was similar for both T-RFLP and Metagenomics methods (Fig. 4B), although the latter showed higher variability. For the metagenomics dataset we also calculated the lower-order Hill numbers (^0^D, ^1^D) which give higher weight to less abundant OTUs. They displayed the same parabolic pattern (Fig. 4C). Welch’s ANOVA tests were statistically significant for all Hill numbers (P < 0.01, P = 0.022 for ^2^D_Metagenomics_). Additionally, there were strong correlations between *α*- diversity and community function (Fig. S6), focusing on the more robust estimators of microbial diversity ^1^D and ^2^D (48). Both ^1^D and ^2^D correlated positively with nitrification and biomass productivity (r > 0.44, P < 0.03), but negatively with organic carbon removal (r < −0.45, P < 0.03).

**FIG 4.**
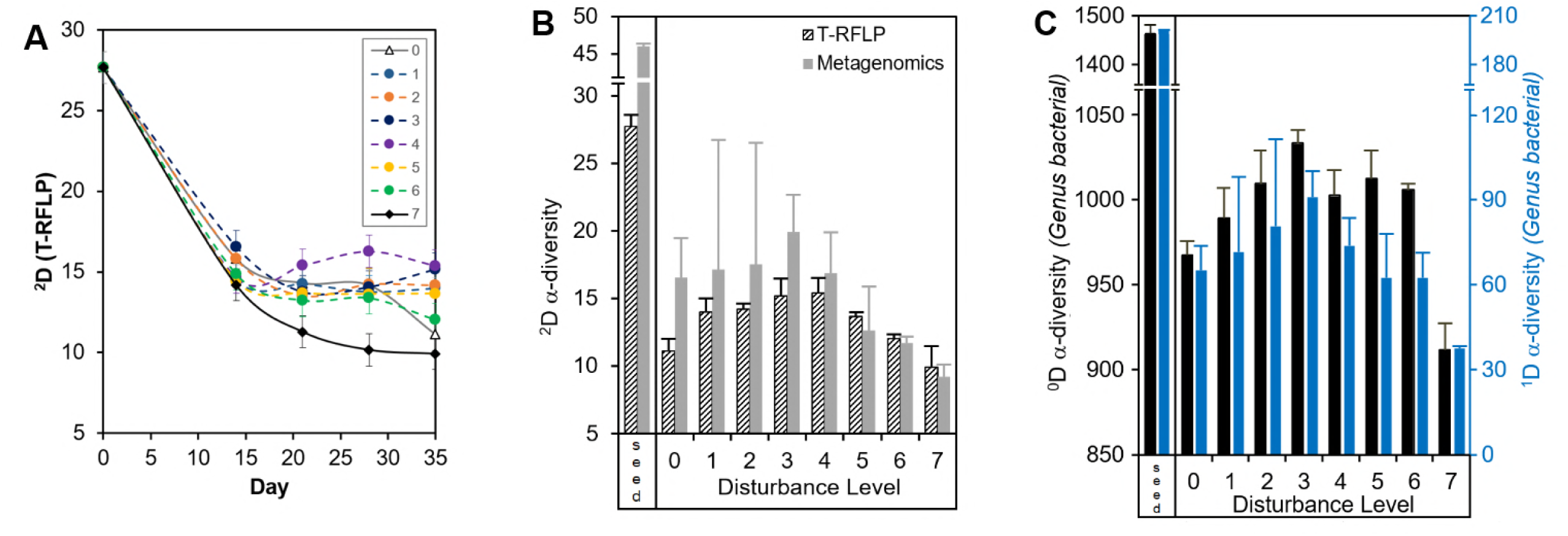
*α*-diversity patterns based on T-RFLP and metagenomics data. (A) Temporal dynamics of Hill number ^2^D for abundant OTUs, calculated from T-RFLP data across disturbance levels. (B) Hill number ^2^D calculated from T-RFLP (black dashed bars) and metagenomics (grey solid bars) data at days 0 (seed) and 35. (C) Hill numbers ^0^D (black solid bars) and ^1^D (blue solid bars) from metagenomics data on days 0 (seed) and 35. Values represent mean ± s.d. (n = 3).

### Null model analyses validate variations in **β**-diversity and temporal increase in stochastic community assembly

To test if the observed changes in **β**-diversity (Fig. 1A, Fig. 3) were due to underlying stochastic and deterministic mechanisms or due to changes in *α*– and *γ*–diversity ratios (*α*:*γ*) alone (49), we employed a null model analysis on the bacterial genus-level metagenomics and T-RFLP datasets on day 35 (Fig. 5A). Linear regression analyses between the calculated *β*-deviation and the observed *γ*-diversity yielded non-significant results (P_Metagenomics_ = 0.42, P_T-RFLP_ = 0.31), confirming that observed changes in *β*-diversity were not only due to changes in *α*:*γ*.

**FIG 5.**
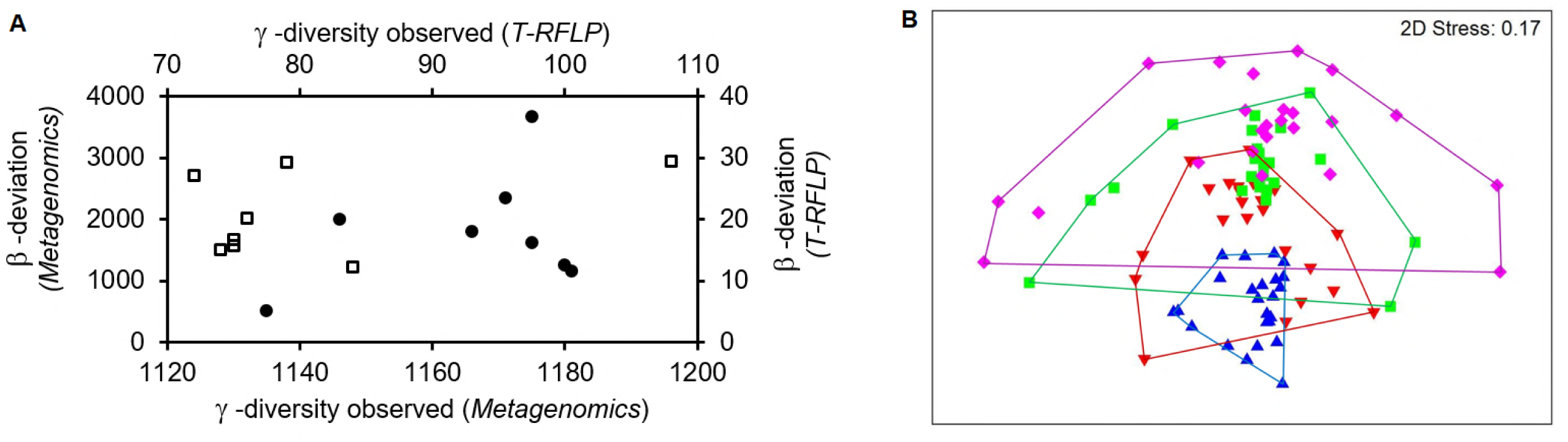
Null model analyses of community data. (A) *β*-deviation versus observed *γ*-diversity for all disturbance treatment levels on day 35. Each point involved all replicates (n = 3) of each disturbance level (a = 8) using the model from (49) on the metagenomics (closed circles, bacterial genus taxonomical level), and T-RFLP (open squares) datasets. Non-significant linear regression tests confirmed that changes in *β*-diversity did not arise from changes in *α*:*γ* diversity alone, suggesting other mechanisms underlying community assembly. (B) NMDS ordination plot of community dynamics of common OTUs (T-RFLP dataset, n = 96) using the Raup-Crick dissimilarity metric as in (51). Days: 14[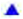], 21[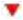], 28[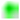], and 35[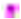]. Lines denote the minimum convex shapes around the data for each disturbance level. Points that are further separated represent communities less deviant from the null expectation, whereas those that are closer together are more divergent from the null expectation. Communities on day 35 are nearer to the null expectation compared with the ones on previous days, suggesting an increasing role over time of stochastic processes in driving community assembly. Overall, null model analyses validate variations in **β**-diversity and temporal increase of stochastic community assembly

Furthermore, another null model analysis was employed to test whether the increase in community dissimilarity observed over time (Fig. 1B, Fig. 3) was related to an increase in stochastic effects driving community assembly. A temporal analysis on the T-RFLP community dataset (n = 96) employing the Raup-Crick dissimilarity metric (50, 51) showed that communities became closer to the null expectation over time, with the greater effect of stochasticity observed after 35 days (Fig. 5B). The observed differential heteroscedasticity among disturbance levels was statistically significant (PERMDISP P = 0.003).

## Discussion

### Deterministic and stochastic patterns of assembly amongst different disturbance levels

Niche-structuring at both ends of the disturbance range was suggested by community assembly patterns and ecosystem function. The undisturbed (L0) and press-disturbed (L7) levels were distinct from each other as well as from the remaining intermediate levels, as supported by multivariate tests (both distance-based and GLMMs). The ordination plots and cluster analyses showed a clear separate clustering for the independent replicates of these two disturbance levels along the experiment, and particularly the constrained ordination plots displayed this with 0% misclassification error. Furthermore, community function was clearly differentiated between L0 and L7, as well as being consistent across replicates at each level. We contend that the observed clustering is an indication that both the undisturbed and press-disturbed levels favoured deterministic assembly mechanisms, where the selective pressure due to unaltered succession (L0) or sustained toxic-stress (L7) promoted species sorting resulting in similar community structuring among biological replicates over the course of the experiment.

Conversely, the communities from intermediately disturbed levels (L1-6) did not form distinct clusters for any particular level through the experiment. Within-treatment dissimilarity among same-level replicates increased over time, with some replicates being more similar to those of other intermediate levels. Concurrently, ecosystem function parameters also displayed within-treatment variability for L1-6. For example, the conversion of ammonia to NO_X_ products, which was initially hampered when communities were still adapting to degrade 3-CA, was not the same across all equally handled independent replicates. The observed divergence across independent replicates is considered here as strong indicator of stochasticity in community assembly. Other aspects might promote stochastic assembly, like strong feedback processes (52) that are linked to density dependence and species interactions (53), priority effects (3), and ecological drift (54). Nonetheless, the observed increasing role of stochasticity over time was also supported by null model analyses, which also validated that the observed changes in *β*-diversity were not due to changes in *α*:*γ* diversity ratios alone (49).

We argue that there were different underlying neutral-niche mechanisms operating in the resulting community assembly along the disturbance range of our study. Similarly, a study on groundwater microbial communities (6) found through null model analysis that both deterministic and stochastic processes played important roles in controlling community assembly and succession, but their relative importance was time dependent. The greater role of stochasticity we found on day 35 concurred with higher observed variability in community function and structure among replicates for intermediately-disturbed levels. Likewise, previous work on freshwater ponds tracking changes in producers and animals (51) found *β*- diversity (in terms of dissimilarity) increasing with stochastic processes. These observed patterns are also in accordance with ecological studies proposing deterministic and stochastic processes balancing each other to allow coexistence (12), with communities exhibiting variations in the strength of stabilization mechanisms and the degree of fitness equivalence among species (11). Thus, it is not sufficient to ask whether communities mirror either stochastic or deterministic processes (10), but also necessary to investigate the combination of such mechanisms that in turn explain the observed community structures along a continuum (11).

### Diversity-disturbance patterns and trade-offs with function

We observed the highest *α*-diversity at intermediate levels as predicted by the IDH (31), both in terms of composition (^0^D) and abundances (^1^D, ^2^D). This finding is non-trivial in two aspects. First, Svensson *et al.* (32) have shown that most studies find support for the IDH when using species richness (^0^D) rather than evenness or other abundance-related indices (like ^1^D and ^2^D). They called for the use of logical arguments to support the idea that peaks in compound diversity-indices should be also expected at intermediate levels of disturbance. Second, the use of richness for microbial communities is not reliable (48) since it is heavily constrained by the method of measurement (55), which makes it hard to compare results from different studies using this metric. Hence, for microbial systems, it is more reasonable to assess diversity in terms of more robust compound indices rather than richness, reason why we focused on ^1^D and ^2^D for diversity-function analyses.

Importantly, the observed pattern in *α*-diversity was time dependent and resulted in an IDH pattern after 35 days. Temporal dynamics were expected since the sludge community experienced an initial perturbation in all reactors after transfer from a wastewater treatment plant to our microcosm arrangement. For the sludge inoculum this implied changes in reactor volume, frequency of feeding (continuous to batch), type of feeding (sewage to complex synthetic media), immigration rates (open to closed system), and mean cell residence time (low to high). This was a succession scenario in which communities had to adapt to such changes along with the designed disturbance array. For L0 and L7, ^2^D decreased over time in agreement with niche-dominated processes, probably because such levels represented the most predictable environments within our disturbance range. In contrast, intermediate levels either increased or maintained the same ^2^D over time (after an initial decrease within the first two weeks), seemingly a case where niche overlap promoted stochastic assembly (10). The emergence of a IDH pattern after time is coherent with findings in previous microcosm studies using synthetic protist (56) and freshwater enrichment (35) microbial communities. Yet, none of these studies evaluated the relative importance of the underlying assembly mechanisms of assembly on the observed diversity dynamics.

Additionally, both ^1^D and ^2^D were positively correlated with nitrification and productivity, suggesting that higher community evenness favours functionality under selective pressure (57), but were negatively correlated with organic carbon removal. Thus, we cannot affirm that more diverse communities have better functionality without considering tradeoffs. This supports the notion that higher *α*-diversity does not necessarily imply a “better” or “healthier” system (55). In addition to the observed OTU diversity, there was an evident ecosystem function diversity along the disturbance range studied, a similar finding to that of previous studies with simpler planktonic communities (58).

Functional tradeoffs are expected under disturbance since organisms need to allocate resources normally used for other functions to recover after a disturbance (59). In our study, communities with higher biomass had lower organic carbon removal efficiencies, which together with the tradeoffs described for nitrification, suggest the adoption of different community life-history strategies depending on the frequency of disturbance. The results presented here were all taxonomy-independent since our focus was on diversity, function, and mechanisms of community assembly (phylum-level community changes are provided as supplemental material Fig. S7). Taxonomy-independent approaches continue to be useful to describe diversity patterns and mechanisms of community assembly (2, 60). However, it has been proposed that species’ traits can predict the effects of disturbance and productivity on diversity (61). Hence, further analysis of the different taxa and their genetic potential paired with the observed tradeoffs in community function will aid in the understanding of potential life-history strategies (59) and their relationship with community aggregated traits (62) in the near future.

### Merging mechanisms of community assembly and alpha-diversity patterns: an intermediate stochasticity hypothesis

Knowing that the validity of the IDH is still under debate (37, 38) and that many different diversity-disturbance patterns have been reported (27, 30, 33), we asked whether there is a relationship between the peaked pattern in diversity observed and the underlying neutral-niche processes of community assembly. Under purely neutral processes, diversity should vary randomly as all species have equal fitness (63), unless some other mechanism acts to prevent this. We hypothesized that higher *α*-diversity at intermediate disturbance frequencies is the result of weaker stabilizing mechanisms (niches), which are stronger at extreme ends of the disturbance range. Neutral (stochastic) mechanisms will produce even assemblages (higher *α*-diversity) at intermediately disturbed levels, whilst infrequent or too-frequent disturbances will favour some species over others (lower *α*-diversity). We propose this idea as the Intermediate Stochasticity Hypothesis (ISH, Fig. 6) and contend that it should hold particularly for compound *α*-diversity indices (48), since the underlying assembly mechanisms would affect taxa abundance distributions.

**FIG 6.**
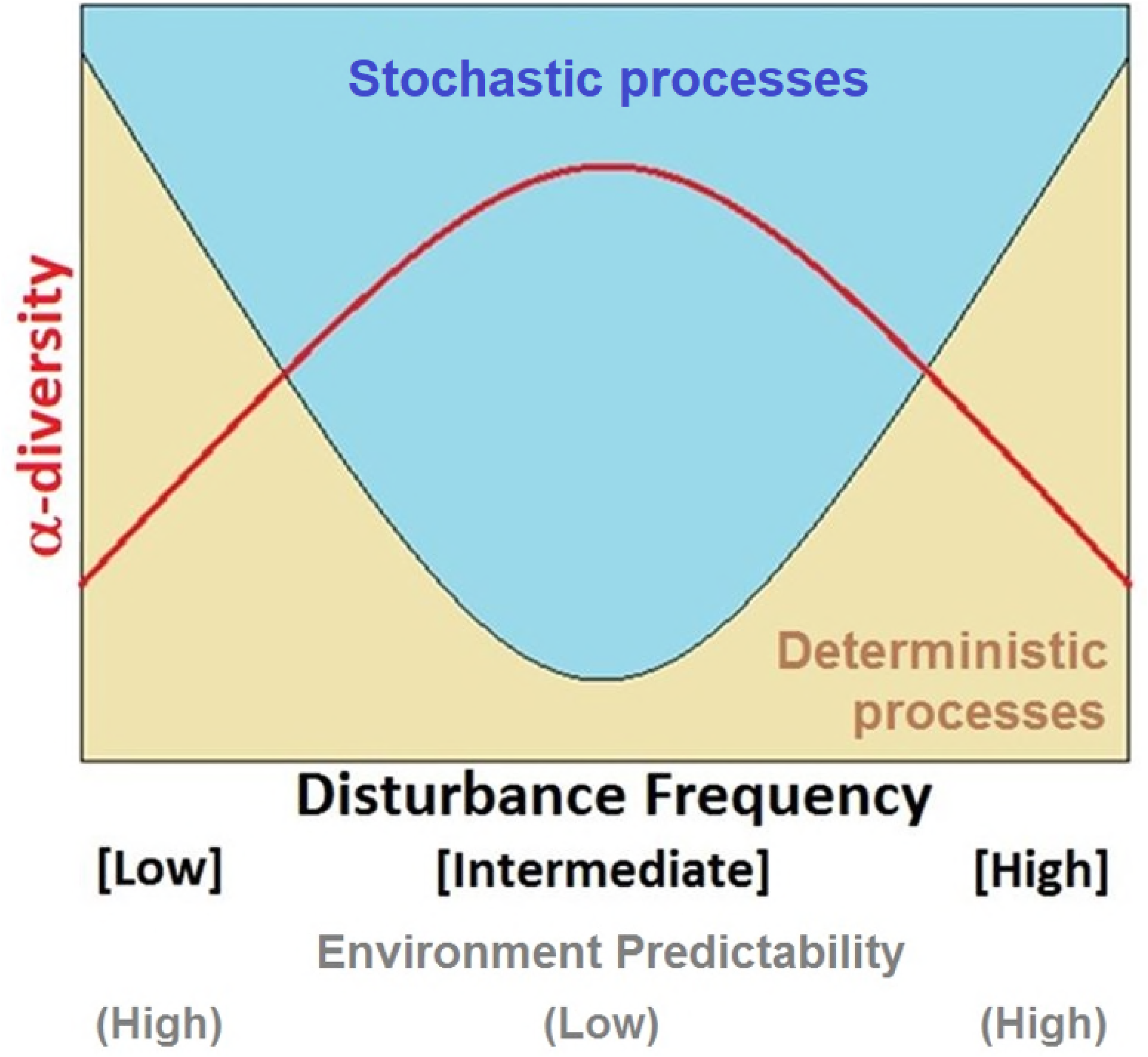
Intermediate Stochasticity Hypothesis (ISH) for community assembly under varying disturbances. Conceptual representation of the classic relationship between *α*-diversity and disturbance (31), including the effect of underlying stochastic and deterministic processes driving bacterial community assembly. When intermediate disturbance regimes result in less predictable environments, specialized traits would be advantageous to taxa, and the stochastic equalization of competitive advantages would lead to higher *α*-diversity. On the contrary, extreme ends of the range where conditions are recurrent would select for adapted organisms whose dominance would result in a lower *α*-diversity.

It is recognized that, beyond empirical pattern description, an understanding of the underlying mechanisms is necessary to comprehend the outcomes of intermediate disturbance regimes (30, 40, 64). Furthermore, microbes constitute a special case since they evolve quickly, compared with plants and animals microbial turnover is rapid, and selection can cause very rapid shifts in community structure (65, 66). The complementary role of stochastic-niche processes could then be one of the key substantive issues promoting species coexistence (37, 39). This can foster research on disturbance-diversity-relationships by complementing the already proposed modifications to the competition-colonization tradeoffs under succession on which the IDH was originally based (31, 38), like considering effects of competitive exclusion (56, 67), productivity (68, 69), spatial heterogeneity (70), feedbacks between diversity and disturbance (71), and type of *α*-diversity metric employed (32).

### Implications and concluding remarks

The implications of this study relate to both process engineering and environmental management. Sludge communities within wastewater treatment are not only model systems in microbial ecology (72), but also a key driver for water sanitation and the environmental impact of anthropogenic water discharges (73). Disturbances could promote stochastic assemblages of the sludge communities, which despite harbouring higher diversity could lead to variable overall community function. This could be the reason why after similar perturbations the process outcomes differ, causing operational problems for water utilities (74). Furthermore, cases where disturbance temporally favours stochastic assembly could lead to a different final community after the perturbation (29), which could compromise the expected ecosystem function. More research is needed to identify such scenarios in practice.

We described how different frequencies of disturbance affected ecosystem function and bacterial community diversity and assembly. Communities were assessed through different molecular methods that nonetheless yielded very similar patterns. Furthermore, besides the wastewater treatment microbial community, other complex microbial systems (e.g. gut microbiome) might display similar responses to disturbance. We argue that changes not only in diversity but also in the underlying deterministic-stochastic assembly mechanisms should be evaluated in studies of the effects of disturbance on such systems. For such an assessment, both replication and wide-enough disturbance ranges are key. This calls for more studies in microcosm (45, 75) and mesocosm settings, as well as meta-analysis from full-scale application studies.

Finally, the ISH could conceptually predict bacterial community shifts in diversity and ecosystem function, given a range of possible disturbance types, in a well-replicated time-series experiment, like it did in this study. Since microbes drive the Earth’s biogeochemical cycles (76), its potential application in biodiversity conservation efforts and the built environment, should encourage further testing under different disturbance dimensions and microbial systems.

## Materials and Methods

### Experimental design

We employed sequencing batch microcosm bioreactors (20-mL working volume) inoculated with activated sludge from a full-scale plant (Text S1) and operated for 35 days. The daily complex synthetic feed included toxic 3-CA at varying frequencies. Eight levels of disturbance were set in triplicate independent reactors (n = 24), which received 3-CA every day (press-disturbed), every two, three, four, five, six, or seven days (intermediately-disturbed), or never (undisturbed). Level numbers were assigned from 0 to 7 (0 for no disturbance, 1 to 7 for low to high disturbance frequency, Fig. S1). Ecosystem function, in the form of process performance parameters, was measured weekly in accordance with Standard Methods (77) where appropriate, and targeted chemical oxygen demand (COD), nitrogen species (ammonium, nitrite, and nitrate ions), volatile suspended solids (VSS), and 3-CA. On the initial day and from the second week onwards, sludge samples (1 mL) were collected weekly for DNA extraction.

### Microbial community analysis and statistical tests

All reported p-values for statistical tests in this study were corrected for multiple comparisons using a False Discovery Rate (FDR) of 10% (78). Community assembly was assessed by a combination of ordination methods (PCO, NMDS, CAP) and multivariate tests (PERMANOVA, PERMDISP) (79) on Bray-Curtis dissimilarity matrixes constructed from square-root transformed normalized abundance data using PRIMER (v.7). Additionally, general linear multivariate models (GLMMs), which deal with mean-variance relationships (80), were employed using the *mvabund* R package (81) fitting the metagenomics dataset to a negative binomial distribution, to ensure that the observed differences among groups were due to disturbance levels and not heteroscedasticity. The 500 most abundant genera (97% of total reads abundances) were employed to ensure random distribution of residuals fitted in the model. Significance was tested using the *anova* function in R with PIT-trap bootstrap resampling (n = 999) (82). Hill diversity indices (83) were employed to measure *α*-diversity as described elsewhere (48, 84), and calculated for normalized non-transformed relative abundance data (Text S1).

Abundance data from Terminal Restriction Fragment Length Polymorphism (T-RFLP) analysis of the bacterial 16S rRNA gene (using 530F-1050R primer set targeting V4-V5 regions) were employed in combination with shotgun metagenomics sequencing at the genus taxonomic level. Such an approach is valid for the questions asked in this study, since comparisons between NGS and fingerprinting techniques support the use of T-RFLP to detect meaningful community assembly patterns and correlations with environmental variables (60), and such patterns can be validated by NGS on a subset of the fingerprinting dataset (2).

### Comparison between Metagenomics and T-RFLP community datasets

Mantel and Procrustes tests (85) were applied to compare metagenomics and T-RFLP datasets from all bioreactors on day 35 (n = 24, subsample of the full T-RFLP dataset). Briefly, Bray-Curtis dissimilarity matrixes were computed using square root transformed T-RFLP data and bacterial genus-level taxa tables generated using a metagenomics approach. Mantel tests were then used to determine the strength and significance of the Pearson product-moment correlation between complete dissimilarity matrices. Procrustes tests (PROTEST) were also employed as an alternative approach to Mantel tests in order to compare and visualize both matrices on PCO and NMDS ordinations. The resultant m2-value is a statistic that describes the degree of concordance between the two matrices evaluated (86). All these statistical tests were performed using the *vegan* R package (functions: *procuste*, *mantel*, *metaMDS*, *vegdist*).

### Metagenomics sequencing and reads processing

Around 173 million paired-end reads were generated in total and 7.2 ± 0.7 million paired-end reads on average per sample. Illumina adaptors, short reads, low quality reads or reads containing any ambiguous base were removed using cutadapt (–m 50 –q 20 - --max-n 0, v.1.11) (87). Taxonomic assignment of metagenomics reads was done following the method described by Ilott *et al.* (88). High quality reads (99.2±0.09% of the raw reads) were randomly subsampled to an even depth of 12,395,400 for each sample prior to further analysis. They were aligned against the NCBI non-redundant (NR) protein database (March 2016) using DIAMOND (v.0.7.10.59) with default parameters (89). The lowest common ancestor approach implemented in MEGAN Community Edition v.6.5.5 (90) was used to assign taxonomy to the NCB-NR aligned reads with the following parameters: maxMatches=25, minScore=50, minSupport=20, paired=true. On average, 48.2% of the high-quality reads were assigned to cellular organisms, from which in turn 98% were assigned to the bacterial domain. Adequacy of sequencing depth was corroborated with rarefaction curves at the genus taxonomy level using MEGAN (90). We did not include genotypic information because it was outside the scope of our study, but will do so in future investigations arising from this work.

### Null Model Analyses on Diversity

*β*-diversity is normally seen as the variation in composition of communities among sites (91), although there are several definitions for it (92). It is also known that most changes in *β*- diversity are determined by variation in the ratio of local (*α*) and regional (*γ*) diversity. Thus, to disentangle the roles of stochastic and deterministic processes as drivers of change in *β* - diversity it is necessary to incorporate statistical null models in the analysis (51, 91), which assume that species interactions are not important for community assembly (93). We employed two null model approaches previously tested in community ecology (49, 51), and more recently for microbial communities (6). To adapt them to handle microbial community data, we considered species as OTUs (i.e. genus for metagenomics and T-RFs for T-RFLP datasets) and each individual count as one read within the metagenomics dataset, or one peak-area unit within the T-RFLP dataset. The first null model approach used here, originally applied to woody plants (49), randomizes the location of each individual within the three independent reactors for each of the eight disturbance treatment levels, while maintaining *γ*- diversity, the total quantity of individuals per reactor, and the relative abundance of each OTU. The second null model uses a modified Raup-Crick dissimilarity metric (*β*_RC_), which indicates the degree to which the observed number of shared OTUs between any two communities deviates from the expected number of shared OTUs in the null model. The closer to the null expectation, the stronger the stochastic effects on the community assembly are, and vice versa (51). This method is robust for variation in *α*-diversity among communities.

## Sequence data and metadata

The microbial DNA metagenomics sequencing datasets supporting the results in this article is available at NCBI BioProjects with accession number: 389377.

## Acknowledgements

This research was supported by the Singapore National Research Foundation and Ministry of Education under the Research Centre of Excellence Program. We thank C.W. Liew for help with sludge sampling, C.F. Liew for technical assistance with the ABI 3730XL DNA analyzer, and S.R. Lohar for the library preparations for metagenomics. F. Lauro, R.B.H. Williams, and S. Kjelleberg are acknowledged for their comments on an earlier version of the manuscript. We thank M. Holyoak for his critical and detailed feedback, as well as E. M. Marzinelli for discussions on data transformations and GLMMs. E.S. was partially supported by a Fulbright fellowship.

## References

1. Stegen JC, Lin XJ, Fredrickson JK, Chen XY, Kennedy DW, Murray CJ, et al. 2013. Quantifying community assembly processes and identifying features that impose them. ISME J 7:2069–2079.

2. Powell JR, Karunaratne S, Campbell CD, Yao H, Robinson L, Singh BK. 2015. Deterministic processes vary during community assembly for ecologically dissimilar taxa. Nat Commun 6:1–10.

3. Zhou J, Ning D. 2017. Stochastic Community Assembly: Does It Matter in Microbial Ecology? Microbiol Mol Biol Rev 81:1–32.

4. Mouillot D, Graham NAJ, Villeger S, Mason NWH, Bellwood DR. 2013. A functional approach reveals community responses to disturbances. Trends Ecol Evol 28:167–177.

5. Briones A, Raskin L. 2003. Diversity and dynamics of microbial communities in engineered environments and their implications for process stability. Curr Opin Biotechnol 14:270–276.

6. Zhou JZ, Deng Y, Zhang P, Xue K, Liang YT, Van Nostrand JD, et al. 2014. Stochasticity, succession, and environmental perturbations in a fluidic ecosystem. Proc Natl Acad Sci USA 111:E836–E845.

7. Griffin JS, Wells GF. 2017. Regional synchrony in full-scale activated sludge bioreactors due to deterministic microbial community assembly. ISME J 11:500–511.

8. Rosindell J, Hubbell SP, Etienne RS. 2011. The Unified Neutral Theory of Biodiversity and Biogeography at Age Ten. Trends Ecol Evol 26:340–348.

9. Silvertown J. 2004. Plant coexistence and the niche. Trends Ecol Evol 19:605–611.

10. Gravel D, Canham CD, Beaudet M, Messier C. 2006. Reconciling niche and neutrality: the continuum hypothesis. Ecol Lett 9:399–409.

11. Adler PB, HilleRisLambers J, Levine JM. 2007. A niche for neutrality. Ecol Lett 10:95–104.

12. Vergnon R, Dulvy NK, Freckleton RP. 2009. Niches versus neutrality: uncovering the drivers of diversity in a species-rich community. Ecol Lett 12:1079–1090.

13. Chase JM, Myers JA. 2011. Disentangling the importance of ecological niches from stochastic processes across scales. Philosophical Transactions of the Royal Society B-Biological Sciences 366:2351–2363.

14. Fisher CK, Mehta P. 2014. The transition between the niche and neutral regimes in ecology. Proceedings of the National Academy of Sciences 111:13111–13116.

15. Ofiteru ID, Lunn M, Curtis TP, Wells GF, Criddle CS, Francis CA, et al. 2010. Combined niche and neutral effects in a microbial wastewater treatment community. Proc Natl Acad Sci USA 107:15345–15350.

16. Jeraldo P, Sipos M, Chia N, Brulc JM, Dhillon AS, Konkel ME, et al. 2012. Quantification of the relative roles of niche and neutral processes in structuring gastrointestinal microbiomes. Proc Natl Acad Sci USA 109:9692–9698.

17. Pholchan MK, Baptista JD, Davenport RJ, Sloan WT, Curtis TP. 2013. Microbial community assembly, theory and rare functions. Front Microbiol 4:1–9.

18. Dini-Andreote F, Stegen JC, van Elsas JD, Salles JF. 2015. Disentangling mechanisms that mediate the balance between stochastic and deterministic processes in microbial succession. Proc Natl Acad Sci USA 112:E1326–E1332.

19. Caruso T, Chan YK, Lacap DC, Lau MCY, McKay CP, Pointing SB. 2011. Stochastic and deterministic processes interact in the assembly of desert microbial communities on a global scale. ISME J 5:1406–1413.

20. Stegen JC, Lin XJ, Konopka AE, Fredrickson JK. 2012. Stochastic and deterministic assembly processes in subsurface microbial communities. ISME J 6:1653–1664.

21. Wang JJ, Shen J, Wu YC, Tu C, Soininen J, Stegen JC, et al. 2013. Phylogenetic beta diversity in bacterial assemblages across ecosystems: deterministic versus stochastic processes. ISME J 7:1310–1321.

22. Dumbrell AJ, Nelson M, Helgason T, Dytham C, Fitter AH. 2010. Relative roles of niche and neutral processes in structuring a soil microbial community. ISME J 4:337–345.

23. Langenheder S, Szekely AJ. 2011. Species sorting and neutral processes are both important during the initial assembly of bacterial communities. ISME J 5:1086–1094.

24. Lee JE, Buckley HL, Etienne RS, Lear G. 2013. Both species sorting and neutral processes drive assembly of bacterial communities in aquatic microcosms. FEMS Microbiol Ecol 86:288–302.

25. Zhou JZ, Liu WZ, Deng Y, Jiang YH, Xue K, He ZL, et al. 2013. Stochastic assembly leads to alternative communities with distinct functions in a bioreactor microbial community. Mbio 4:1–8.

26. Cain M, Bowman W, Hacker S. Ecology. 3 ed. Sunderland, Massachusetts: Sinauer Associates Inc.; 2014.

27. Mackey RL, Currie DJ. 2001. The diversity-disturbance relationship: Is it generally strong and peaked? Ecology 82:3479–3492.

28. Shade A, Read JS, Youngblut ND, Fierer N, Knight R, Kratz TK, et al. 2012. Lake microbial communities are resilient after a whole-ecosystem disturbance. ISME J 6:2153–2167.

29. Shade A, Peter H, Allison SD, Baho DL, Berga M, Burgmann H, et al. 2012. Fundamentals of microbial community resistance and resilience. Front Microbiol 3:1–19.

30. Miller AD, Roxburgh SH, Shea K. 2011. How frequency and intensity shape diversity-disturbance relationships. Proc Natl Acad Sci USA 108:5643–5648.

31. Connell JH. 1978. Diversity in tropical rain forests and coral reefs. Science 199:1302–1310.

32. Svensson JR, Lindegarth M, Jonsson PR, Pavia H. 2012. Disturbance–diversity models: what do they really predict and how are they tested? Proceedings of the Royal Society B: Biological Sciences 279:2163–2170.

33. Kershaw HM, Mallik AU. 2013. Predicting Plant Diversity Response to Disturbance: Applicability of the Intermediate Disturbance Hypothesis and Mass Ratio Hypothesis. Crit Rev Plant Sci 32:383–395.

34. Kim M, Heo E, Kang H, Adams J. 2013. Changes in Soil Bacterial Community Structure with Increasing Disturbance Frequency. Microb Ecol 66:171–181.

35. Gibbons SM, Scholz M, Hutchison AL, Dinner AR, Gilbert JA, Coleman ML. 2016. Disturbance Regimes Predictably Alter Diversity in an Ecologically Complex Bacterial System. mBio 7:1–10.

36. Roxburgh SH, Shea K, Wilson JB. 2004. The intermediate disturbance hypothesis: Patch dynamics and mechanisms of species coexistence. Ecology 85:359–371.

37. Fox JW. 2013. The intermediate disturbance hypothesis should be abandoned. Trends Ecol Evol 28:86–92.

38. Sheil D, Burslem D. 2013. Defining and defending Connell’s intermediate disturbance hypothesis: a response to Fox. Trends Ecol Evol 28:571–572.

39. Fox JW. 2013. The intermediate disturbance hypothesis is broadly defined, substantive issues are key: a reply to Sheil and Burslem. Trends Ecol Evol 28:572–573.

40. Shea K, Roxburgh SH, Rauschert ESJ. 2004. Moving from pattern to process: coexistence mechanisms under intermediate disturbance regimes. Ecol Lett 7:491–508.

41. Krol JE, Penrod JT, McCaslin H, Rogers LM, Yano H, Stancik AD, et al. 2012. Role of IncP-1 beta plasmids pWDL7::rfp and pNB8c in chloroaniline catabolism as determined by genomic and functional analyses. Appl Environ Microbiol 78:828–838.

42. Falk MW, Wuertz S. 2010. Effects of the toxin 3-chloroaniline at low concentrations on microbial community dynamics and membrane bioreactor performance. Water Res 44:5109–5115.

43. Jessup CM, Kassen R, Forde SE, Kerr B, Buckling A, Rainey PB, et al. 2004. Big questions, small worlds: microbial model systems in ecology. Trends Ecol Evol 19:189–197.

44. Benton TG, Solan M, Travis JMJ, Sait SM. 2007. Microcosm experiments can inform global ecological problems. Trends Ecol Evol 22:516–521.

45. Drake JM, Kramer AM. 2012. Mechanistic analogy: how microcosms explain nature. Theor Ecol 5:433–444.

46. Prosser JI. 2010. Replicate or lie. Environ Microbiol 12:1806–1810.

47. Anderson MJ, Willis TJ. 2003. Canonical analysis of principal coordinates: A useful method of constrained ordination for ecology. Ecology 84:511–525.

48. Haegeman B, Hamelin J, Moriarty J, Neal P, Dushoff J, Weitz JS. 2013. Robust estimation of microbial diversity in theory and in practice. ISME J 7:1092–1101.

49. Kraft NJB, Comita LS, Chase JM, Sanders NJ, Swenson NG, Crist TO, et al. 2011. Disentangling the Drivers of *β* Diversity Along Latitudinal and Elevational Gradients. Science 333:1755–1758.

50. Chase JM, Kraft NJB, Smith KG, Vellend M, Inouye BD. 2011. Using null models to disentangle variation in community dissimilarity from variation in α-diversity. Ecosphere 2:1–11.

51. Chase JM. 2010. Stochastic Community Assembly Causes Higher Biodiversity in More Productive Environments. Science 328:1388–1391.

52. Soto-Ortiz L. 2015. The Regulation of Ecological Communities Through Feedback Loops: A Review. Research in Zoology 5:1–15.

53. Holyoak M, Loreau M. 2006. Reconciling empirical ecology with neutral community models. Ecology 87:1370–1377.

54. Nemergut DR, Schmidt SK, Fukami T, O’Neill SP, Bilinski TM, Stanish LF, et al. 2013. Patterns and Processes of Microbial Community Assembly. Microbiol Mol Biol Rev 77:342–356.

55. Shade A. 2017. Diversity is the question, not the answer. ISME J 11:1–6.

56. Violle C, Pu ZC, Jiang L. 2010. Experimental demonstration of the importance of competition under disturbance. Proc Natl Acad Sci USA 107:12925–12929.

57. Wittebolle L, Marzorati M, Clement L, Balloi A, Daffonchio D, Heylen K, et al. 2009. Initial community evenness favours functionality under selective stress. Nature 458:623–626.

58. Weithoff G, Walz N, Gaedke U. 2001. The intermediate, disturbance hypothesis - species diversity or functional diversity? J Plankton Res 23:1147–1155.

59. Krause S, Le Roux X, Niklaus PA, Van Bodegom PM, Lennon JT, Bertilsson S, et al. 2014. Trait-based approaches for understanding microbial biodiversity and ecosystem functioning. Front Microbiol 5:1–10.

60. van Dorst J, Bissett A, Palmer AS, Brown M, Snape I, Stark JS, et al. 2014. Community fingerprinting in a sequencing world. FEMS Microbiol Ecol 89:316–330.

61. Haddad NM, Holyoak M, Mata TM, Davies KF, Melbourne BA, Preston K. 2008. Species’ traits predict the effects of disturbance and productivity on diversity. Ecol Lett 11:348–356.

62. Fierer N, Barberan A, Laughlin DC. 2014. Seeing the forest for the genes: using metagenomics to infer the aggregated traits of microbial communities. Front Microbiol 5:1–6.

63. Hubbell SP. The unified neutral theory of biodiversity and biogeography. Monographs in Population Biology. 322001. p. i-xiv, 1-375.

64. Huston MA. 2014. Disturbance, productivity, and species diversity: empiricism vs. logic in ecological theory. Ecology 95:2382–2396.

65. Prosser JI, Bohannan BJM, Curtis TP, Ellis RJ, Firestone MK, Freckleton RP, et al. 2007. The role of ecological theory in microbial ecology. Nat Rev Microbiol 5:384–392.

66. Hanson CA, Fuhrman JA, Horner-Devine MC, Martiny JBH. 2012. Beyond biogeographic patterns: processes shaping the microbial landscape. Nat Rev Microbiol 10:497–506.

67. Huston M. 1979. A General Hypothesis of Species Diversity. The American Naturalist 113:81–101.

68. Kassen R, Buckling A, Bell G, Rainey PB. 2000. Diversity peaks at intermediate productivity in a laboratory microcosm. Nature 406:508–512.

69. Kondoh M. 2001. Unifying the relationships of species richness to productivity and disturbance. Proceedings of the Royal Society B: Biological Sciences 268:269–271.

70. Buckling A, Kassen R, Bell G, Rainey PB. 2000. Disturbance and diversity in experimental microcosms. Nature 408:961–964.

71. Hughes AR, Byrnes JE, Kimbro DL, Stachowicz JJ. 2007. Reciprocal relationships and potential feedbacks between biodiversity and disturbance. Ecol Lett 10:849–864.

72. Daims H, Taylor MW, Wagner M. 2006. Wastewater treatment: a model system for microbial ecology. Trends Biotechnol 24:483–489.

73. Tchobanoglous GB, Franklin L, Stensel HD. Wastewater Engineering: Treatment and Reuse 4ed. Boston: McGraw Hill; 2003.

74. Pholchan MK, Baptista JD, Davenport RJ, Curtis TP. 2010. Systematic study of the effect of operating variables on reactor performance and microbial diversity in laboratory-scale activated sludge reactors. Water Res 44:1341–1352.

75. Altermatt F, Fronhofer EA, Garnier A, Giometto A, Hammes F, Klecka J, et al. 2015. Big answers from small worlds: a user’s guide for protist microcosms as a model system in ecology and evolution. Methods in Ecology and Evolution 6:218–231.

76. Widder S, Allen RJ, Pfeiffer T, Curtis TP, Wiuf C, Sloan WT, et al. 2016. Challenges in microbial ecology: building predictive understanding of community function and dynamics. ISME J 10:2557–2568.

77. APHA-AWWA-WEF. Standard Methods for the Examination of Water and Wastewater. 22 ed. Washington D.C., USA: APHA-AWWA-WEF; 2005.

78. Benjamini Y, Hochberg Y. 1995. Controlling the False Discovery Rate: A Practical and Powerful Approach to Multiple Testing. Journal of the Royal Statistical Society Series B (Methodological) 57:289–300.

79. Clarke KR, Gorley RN. PRIMER v7: User Manual/Tutorial. Plymouth, UK: PRIMER-E; 2015. 296 p.

80. Warton DI, Wright ST, Wang Y. 2012. Distance-based multivariate analyses confound location and dispersion effects. Methods in Ecology and Evolution 3:89–101.

81. Wang Y, Naumann U, Wright ST, Warton DI. 2012. mvabund– an R package for model-based analysis of multivariate abundance data. Methods in Ecology and Evolution 3:471–474.

82. Warton DI, Thibaut L, Wang YA. 2017. The PIT-trap—A “model-free” bootstrap procedure for inference about regression models with discrete, multivariate responses. PLOS ONE 12:e0181790.

83. Hill MO. 1973. Diversity and evenness: a unifiying notation and its consequences. Ecology 54:427–432.

84. Vuono DC, Benecke J, Henkel J, Navidi WC, Cath TY, Munakata-Marr J, et al. 2015. Disturbance and temporal partitioning of the activated sludge metacommunity. ISME J 9:425–435.

85. Peres-Neto PR, Jackson DA. 2001. How well do multivariate data sets match? The advantages of a Procrustean superimposition approach over the Mantel test. Oecologia 129:169–178.

86. Jackson DA. 1995. PROTEST: A PROcrustean Randomization TEST of community environment concordance. Ecoscience 2:297–303.

87. Chen C, Khaleel SS, Huang H, Wu CH. 2014. Software for pre-processing Illumina next-generation sequencing short read sequences. Source Code for Biology and Medicine 9:8–8.

88. Ilott NE, Bollrath J, Danne C, Schiering C, Shale M, Adelmann K, et al. 2016. Defining the microbial transcriptional response to colitis through integrated host and microbiome profiling. ISME J:2389–2404.

89. Buchfink B, Xie C, Huson DH. 2015. Fast and sensitive protein alignment using DIAMOND. Nat Meth 12:59–60.

90. Huson DH, Beier S, Flade I, Górska A, El-Hadidi M, Mitra S, et al. 2016. MEGAN Community Edition - Interactive Exploration and Analysis of Large-Scale Microbiome Sequencing Data. PLoS Comp Biol 12:e1004957.

91. Anderson MJ, Crist TO, Chase JM, Vellend M, Inouye BD, Freestone AL, et al. 2011. Navigating the multiple meanings of beta diversity: a roadmap for the practicing ecologist. Ecol Lett 14:19–28.

92. Tuomisto H. 2010. A diversity of beta diversities: straightening up a concept gone awry. Part 1. Defining beta diversity as a function of alpha and gamma diversity. Ecography 33:2–22.

93. Gotelli NJ, McGill BJ. 2006. Null Versus Neutral Models: What’s The Difference? Ecography 29:793–800.

